# Mutating novel interaction sites in NRP1 reduces SARS-CoV-2 spike protein internalization

**DOI:** 10.1101/2021.10.11.463689

**Authors:** Debjani Pal, Kuntal De, Timothy B. Yates, Wellington Muchero

## Abstract

The global pandemic of Coronavirus disease 2019 caused by severe acute respiratory syndrome coronavirus 2 has become a severe global health problem because of its rapid spread(*1*). Both angiotensin-converting enzyme 2 and neuropilin 1 provide initial viral binding sites for SARS-CoV-2 (*2, 3*). Here, we show that three cysteine residues located in a1/a2 and b1 domains of neuropilin 1 are necessary for SARS-CoV-2 spike protein internalization in human cells. Mutating cysteines C82, C104 and C147 altered neuropilin 1 stability and binding ability as well as cellular internalization and lysosomal translocation of the spike protein. This resulted in up to 4 times reduction in spike protein load in cells for the original, alpha and delta SARS-CoV-2 variants even in the presence of the endogenous angiotensin-converting enzyme 2 receptor. Transcriptome analysis of cells transfected with mutated NRP1 revealed significantly reduced expression of genes involved in viral infection and replication, including eight members of the ribosomal protein L, ten members of ribosomal protein S and five members of the proteasome β subunit family proteins. We also observed higher expression of genes involved in suppression of inflammation and endoplasmic reticulum associated degradation. These observations suggest that these cysteines offer viable targets for therapies against COVID-19.

## Main Text

Neuropilin (NRP) is a single membrane-spanning, type-I transmembrane, non-tyrosine kinase receptor protein(*4*). In humans, it has two homologs, NRP1 and NRP2, with a 44% of sequence identity (*5, 6*). Both contain five extracellular domains which are, two CUB (a1/a2), two coagulation factor FV/FVIII (b1/b2), a MAM (meprin, A5, and μ-phosphatase; also known as c) and a domain that contains a transmembrane and a short cytoplasmic region (*7, 8*). NRP1 was identified as a co-receptor for several extracellular ligands, including class III semaphorins, a specific isoform of the vascular endothelial growth factor-A (VEGF-A), heparin-binding proteins, placental growth factor 2 and fibroblast growth factor 2 (*9, 10*). Both CUB and FV/FVIII domains provide the binding sites for secreted-semaphorins and VEGF (*11*). NRP1 plays an essential role in performing key physiological processes like angiogenesis and semaphorin dependent axon guidance signaling (*12, 13*).

An interesting aspect of NRP1 is that its ligand, semaphorins, are synthesized as inactive precursor forms and require proteolytic cleavage by furin or related endoproteolytic pro-protein convertases, a key feature that is also utilized by the viral coat proteins (*14*). For example, NRP1 has been reported to function as an entry receptor for Epstein-Barr virus (EPV) and more recently for the SARS-CoV-2 virus (*3, 15, 16*). To promote NRP1-mediated cell internalization via tissue penetration, cleaved peptides utilize the R/K/XXR/K motif at their C-terminus (C-end rule) (*17*). CendR peptides are known to be useful for cancer drug delivery, as NRP1 is frequently overexpressed in various tumor types (*18*). This similar phenomenon used by SARS-CoV-2 while binding with NRP1 for virus-host membrane fusion via the multibasic site at the S1-S2 boundary of the viral spike protein (S) (*3*). S-protein usually gets activated by cellular proteases, either by furin and or TMPRSS2, which ensure cleavage at the S1/S2 site (*19*). The S1 subunit of the S protein of coronaviruses facilitates viral entry into target cells by providing a receptor-binding domain (*19*). Since the alarming spread of the COVID-19 pandemic, a major focus of ongoing research has been to define the mechanism behind viral entry. For SARS-CoV-2, the CendR pocket is usually present within the extracellular b1b2 domain of NRP1 (*3*). However, the dynamics of S-protein binding to the NRP1 receptor at the specific amino acid level are yet to be established.

## Results

We previously identified orthologs of NRP1 and NRP2 that carry a Plasminogen-Apple-Nematode (PAN) domain. A key feature of the PAN domain are 4-6 strictly conserved cysteine residues that form its core. This protein domain was implicated in protein ubiquitination and proteolysis of two unrelated proteins, a hepatocyte growth factor in human and a G-type lectin receptor-like kinase in plants(*20*) (De et al. 2021-manuscript under review). Mutation of amino acids in the PAN domain of both proteins led to dramatic changes in immune signaling in both organisms(*20*) (De et al. 2021-manuscript under review). Since the human NRP1 does not have a reported PAN domain, we sought to evaluate if any features of the domain could be detected by assessing residual homology with NRP1 and 2 from other organisms. To do this, we aligned the human NRP1 with a *Macrostomum lignano* NRP2 which carries an intact PAN domain. Based on this analysis, we identified a region of NRP1 that exhibited weak homology to the *M. ligano* PAN domain. Specifically, we identified four cysteine residues at amino acid positions C82, C104, C147 and C173 in a1/a2 and b1/b2 domains of NRP1 which could suggest existence of a vestigial PAN domain (Fig. 1a). We, therefore hypothesized that these residues may play a role in cellular response to viral infection.

**Fig. 1:**
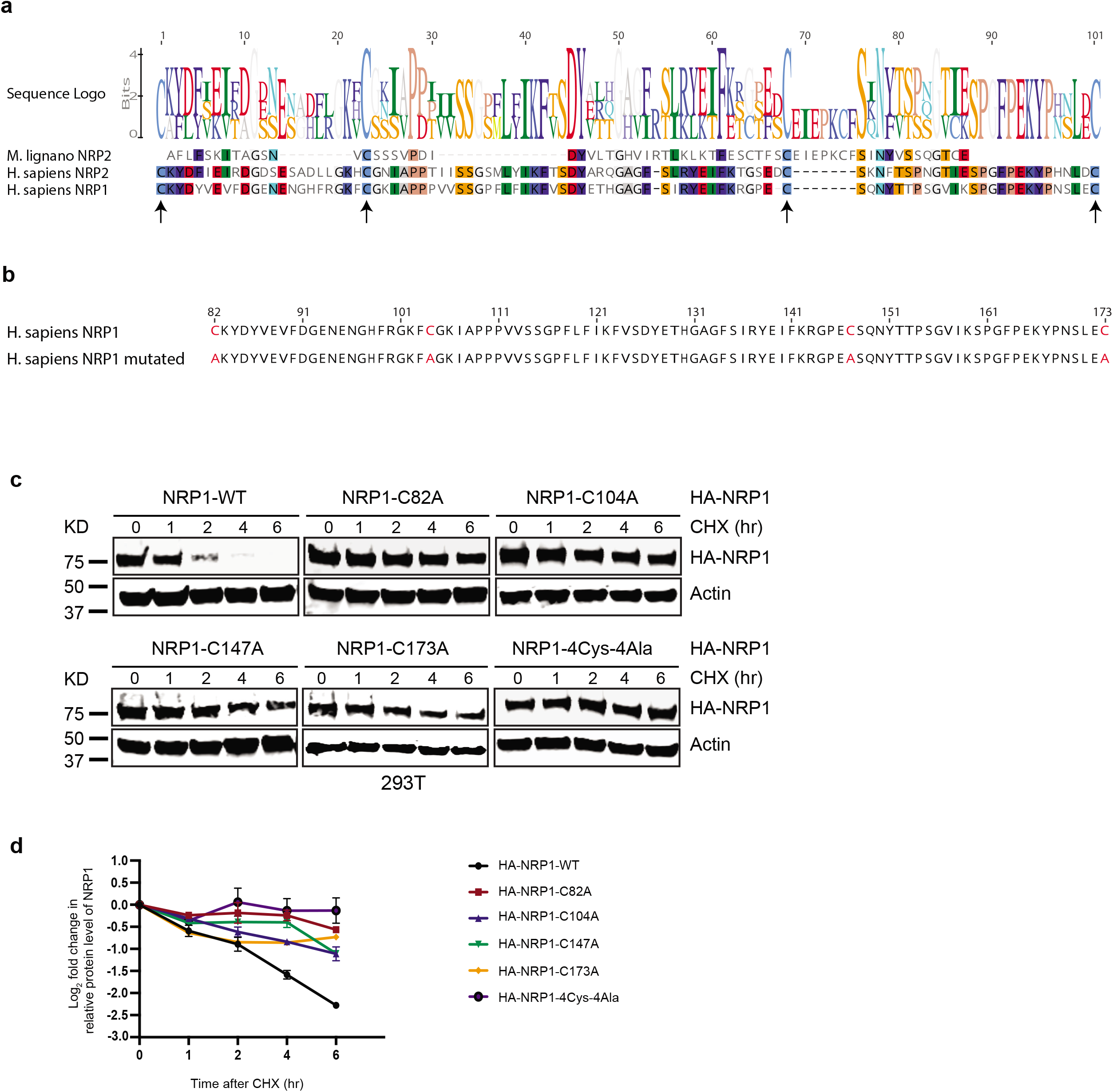
Novel cysteine controls NRP1 stability. (a) Multiple alignment of the sequences of PAN domain of representative proteins from two different organisms highlights the position of four conserved cysteines. (b) Schematic diagram of amino acid sequence represents NRP1 vestigial PAN domain along with the four marked cysteines (Cys82, Cys104, Cys147 and Cys173) and the subsequent mutant version where conserved cysteines were mutated to alanine (Ala82, Ala104, Ala147 and Ala173). (c) Immunoblot analysis of whole cell lysates derived from 293T cells, transfected with HA-NRP1 WT and different single cysteine mutants of HA-NRP1 and HA-NRP1-4Cys-4Ala constructs as indicated. 30 hours post-transfection, whole-cell lysates were prepared for immunoblot analysis. Representative image of n=3 biological replicates. (d) Quantification of the band intensities in (c). The intensities of HA-NRP1 (WT and mutants) bands were normalized to actin and then normalized to HA-NRP1 WT. Data are represented as mean ± SD, n = 3, and *p<0.05, **p < 0.005, *** p < 0.0005 were calculated with one-way ANOVA.

To evaluate the functional significance of these four cysteines, we introduced both single cysteine to alanine mutations as well as mutating all four cysteines to alanines in NRP1 (Fig. 1b). Like most other transmembrane receptors, upon binding with its ligand, NRP1 undergoes internalization and is targeted for lysosomal degradation(*21*). Based on this established model, we first evaluated the overall abundance of cysteine-to-alanine NRP1 proteins in cells compared to the wild type. Based on the cycloheximide (CHX) protein stability assay, we observed that mutations in any of the four-cysteines increased protein stability of exogenously expressed HA-tagged NRP1 in HEK293T cells (Fig. 1c and Fig. 1d). The HA-NRP1 4Cys-4Ala mutant showed a cumulative impact on the protein stability by all the four-cysteines mutation as expected (Fig. 1d). These data suggested that all four cysteines could be involved in targeting of NRP1 protein by degradation pathways in cells.

Secondly, we sought to evaluate impact of these mutations on NRP1 interaction with SARS-CoV-2 S1 protein. SARS-CoV-2 S1 protein residues ^493-685^ have been reported to bind with NRP1(*3*). We made a Flag-SARS-CoV-2-S1^493-685^ construct based on the sequence available in NCBI (Fig. 2a). To assess the impact of these four novel cysteines in NRP1 interaction with COVID-19, we carried out a direct coimmunoprecipitation assay between HA-NRP1-WT, HA-NRP1-C82A, HA-NRP1-C104A, HA-NRP1-C147, HA-NRP1-C173A, HA-NRP1-4Cys-4Ala, and Flag-SARS-CoV-2-S1^493-685^ following expression in HEK293T cells. Mutations in any of the first three cysteines (C82A, C104A, and C147A) in NRP1 had a significantly negative impact on Flag-SARS-CoV-2-S1^493-685^ protein binding as evidenced by quantification of the relative amount of Flag-SARS-CoV-2-S1^493-685^ pulled down with different variants of NRP1 (Fig. 2b and Fig. 2c).

**Fig. 2:**
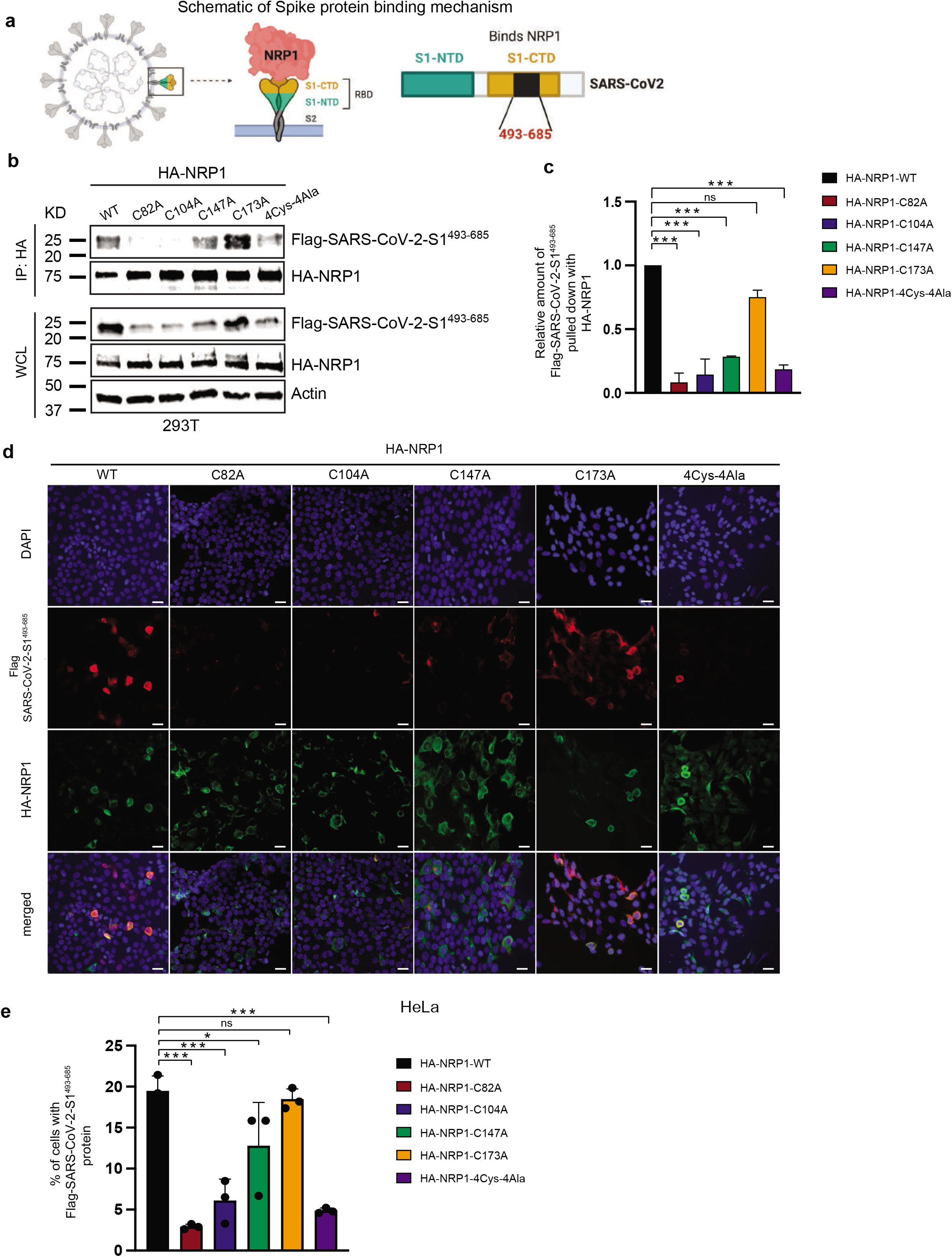
Novel cysteines in NRP1 facilitates its interaction with SARS-CoV-2 spike protein. (a) Schematic representation of SARS-CoV-2 spike protein binding with NRP1 receptor. Contacting residues on spike protein is from 493-685. (b) NRP1 binding with spike protein is novel cysteines dependent. 293T cells were transfected with the indicated HA-NRP1 (wild type and mutants) and Flag-SARS-CoV-2-S1^493-685^ constructs. Cells were lysed 30 hr post transfection, and the interaction between HA-NRP1 and Flag-SARS-CoV-2-S1^493-685^ was analyzed. (c) Quantification of the band intensities (*n* = 2). Immunoprecipitated Flag-SARS-CoV-2-S1^493-685^ band intensities were normalized to the respective HA-NRP1 IP bands and then further normalized to HA-NRP1-WT control. Data are represented as mean ± SD, and *p<0.05, **p < 0.005, *** p < 0.0005 (Student’s t test). (d) Representative images of colocalization studies between Flag-SARS-CoV-2-S1^493-685^ and different HA-NRP1 constructs by confocal immunofluorescence microscopy in HeLa cells. The cells were transiently transfected with Flag-SARS-CoV-2-S1^493-685^ and different mutants of HA-NRP1 as indicated. 30 hours post-transfection cells were fixed, mounted and protein expression patterns were visualized using a Zeiss LSM 710 confocal microscope outfitted with a 63x objective. Scale bars represent 20 μm. The images shown are representative from three independent biological experiments (average 100 cells were observed per experimental condition per replicate). (e) Quantification of the HeLa cells expressing Flag-SARS-CoV-2-S1^493-685^ in the presence of indicated HA-NRP1 constructs. Data are represented as mean ± SD, n = 3 (average 100 cells were observed for each condition per experiment), and *p<0.05, **p < 0.005, *** p < 0.0005 were calculated by one-way ANOVA.

Likewise, immunofluorescence data corroborated these findings and showed notably reduced colocalization between HA-NRP1-C82A, HA-NRP1-C104A, HA-NRP1-C147A, HA-NRP1-4Cys-4Ala and Flag-SARS-CoV-2-S1^493-685^ compared to HA-NRP1-WT (Fig. 2d). Remarkably, the overall viral spike protein (Flag-SARS-CoV-2-S1^493-685^) density was reduced up to four times in cells co-transfected with HA-NRP1-C82A, C104A and HA-NRP1-4Cys-4Ala mutants compared to wild type NRP1 (Fig. 2d and Fig. 2e).

We also tested whether these cysteines facilitated binding of NRP1 with the co-receptor Plexin-A1 as Plexin-A1 helps to form a functional receptor for semaphorin 3A by directly interacting with NRP1(*22*). HA-NRP1-WT and HA-NRP1-4Cys-4Ala variants were co-expressed with Flag-Plexin-A1 in HEK293T cells, and complexes were immunoprecipitated using anti-Flag. Our results indicated that the 4Cys-4Ala mutant was unable to associate with Plexin-A1 (Fig. S1a and S1b). In agreement with our biochemical data, we reconfirmed that mutating these cysteine residues reduced the colocalization signal for NRP1 and Plexin-A1 when transfected in HeLa cells (Fig. S1c).

We then examined whether full-length SARS-CoV-2-S protein could influence the interaction with NRP1. Fig. 3a shows that full-length SARS-CoV-2-S protein is equally susceptible to NRP1 cysteine mutations. The interaction between HA-NRP1-4Cys-4Ala and SARS-CoV-2-S was significantly impaired (Fig. 3b). The immediate development of a vaccine against COVID-19 has provided a vital tool to fight against the rapid spread of the SARS-CoV-2 virus. However, constantly evolving mutations in the S protein imposes a challenge for effective deployment of vaccines.

**Fig. 3:**
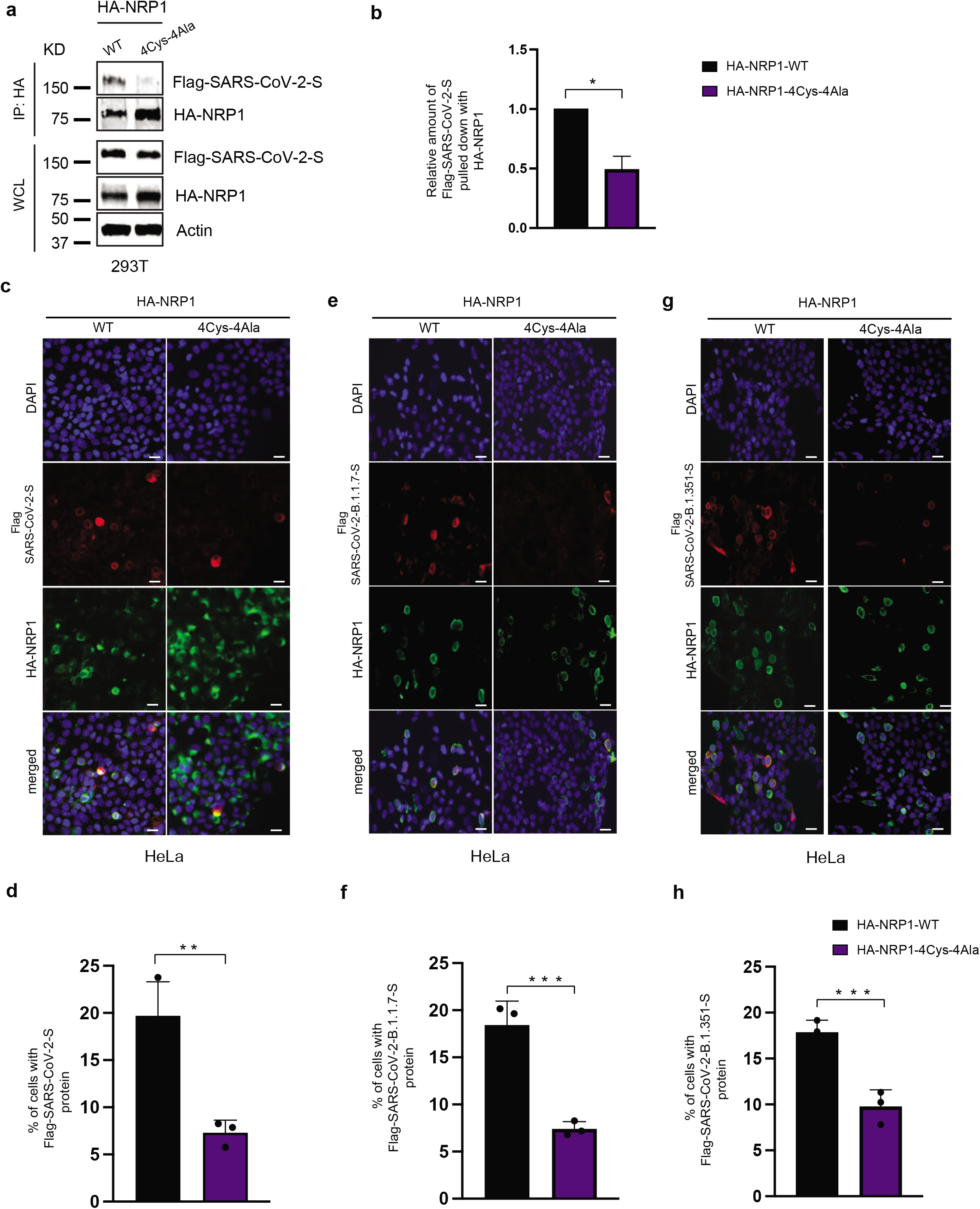
Mutation in the novel cysteines in NRP1 universally impairs its interaction with different SARS-CoV-2 spike protein variants. (a) NRP1 PAN domain cysteines play crucial role in SARS-CoV-2 spike protein binding. 293T cells were transfected with the indicated HA-NRP1 (wild type and 4Cys-4Ala mutant) and Flag-SARS-CoV-2-S constructs. Cells were lysed 30 hr post transfection, and the interaction between HA-NRP1 and Flag-SARS-CoV-2-S was analyzed. (b) Quantification of the band intensities (*n* = 3). Immunoprecipitated Flag-SARS-CoV-2-S band intensities were normalized to the respective HA-NRP1 IP bands and then further normalized to HA-NRP1-WT control. Data are represented as mean ± SD, and *p<0.05, **p < 0.005, *** p < 0.0005 (one-way ANOVA). (c, e, g) Representative images of colocalization studies between Flag-tagged different SARS-CoV-2 spike protein variants and indicated HA-NRP1 constructs by confocal immunofluorescence microscopy in HeLa cells. The cells were transiently transfected with different Flag-SARS-CoV-2-S and different constructs of HA-NRP1 as indicated. 30 hours post-transfection cells were fixed, mounted and protein expression patterns were visualized using a Zeiss LSM 710 confocal microscope outfitted with a 63x objective. Scale bars represent 20 μm. The images shown are representative from three independent biological experiments (average 100 cells were observed per experimental condition per replicate). (d,f,h) Quantification of the HeLa cells expressing Flag-SARS-CoV-2-S, Flag-SARS-CoV-2-B.1.1.7-S and Flag-SARS-CoV-2-B.1351-S in the presence of indicated HA-NRP1 constructs. Percentage of cells with Flag-SARS-CoV-2-S protein were calculated for each of the variant separately and normalized compared to their respective control set (cells with HA-NRP1-WT). Data are represented as mean ± SD, n = 3 (average 100 cells were observed for each condition per experiment), and *p<0.05, **p < 0.005, *** p < 0.0005 were calculated by one-way ANOVA.

To establish NRP1 as a potential universal candidate for targeted therapy, we used full-length Flag-tagged SARS-CoV-2-S, SARS-CoV-2-B.1.1.7-S and Flag-tagged SARS-CoV-2-B1.351-S, representing the original, alpha and beta variants, respectively and used those for colocalization assays. HA-NRP1-4Cys-4Ala was equally incapable of interacting with the variants, as confirmed from the immunofluorescence data (Fig. 3c, e, and g). Additionally, viral spike protein load in cells was reduced significantly (Fig. 3d, f, and h). This study provides the first experimental evidence supporting our hypothesis that targeting these four specific amino acids in NRP1 could be effective in reducing viral spread irrespective of the spike protein variants, although emerging variants will need to be evaluated.

SARS-CoV-2 follow the conventional endosomal secretory route for its subsequent maturation and release from the infected cells(*23*). However, Ghosh et al. recently described that mouse hepatitis virus (MHV, member of β-coronavirus family like SARS-CoV-2) use lysosomal Arl8b-dependent pathway for the egress(*24*). Late endosomes/lysosome deacidification inhibits lysosomal proteases and favor the egress(*24*). We investigated whether transfected SARS-CoV-2 spike protein showed any difference in lysosomal association when co-expressed with 4Cys-4Ala mutant NRP1 in cells. We transfected HeLa cells with Flag tagged SARS-CoV-2 S and HA tagged WT and 4Cys-4Ala NRP1s then co-stained with Flag and the late endosomal/lysosomal transmembrane protein LAMP1 (Fig. S2). Interestingly, there was significant reduction in the LAMP1 association of spike protein when transfected with NRP1-4Cys-4Ala and observed under confocal microscopy.

To ascertain that our observations were related to lower spike protein density in cells, we co-transfected HA-NRP1-WT and NRP1-4Cys-4Ala mutant with Flag-SARS-CoV-2-S1^493-685^ at different ratios and measured protein levels (Fig. 4a). Quantification from 3 biological replicates further supported our previous observation that reduced spike protein density was solely due to the lack of functional NRP1 receptor in cells (Fig. 4b). In fact, increasing the transfection ratio for Flag-SARS-CoV-2-S1^493-685^ could not fully rescue the spike protein yield in cells transfected with HA-NRP1-4Cys-4Ala mutant (Fig. 4a and 4b). Full-length SARS-CoV-2-S protein also exhibited reduced spike protein density when co-transfected in a 1:1 ratio together with the HA-NRP1-4Cys-4Ala mutant compared to wild type (Fig. 4c).

**Fig. 4:**
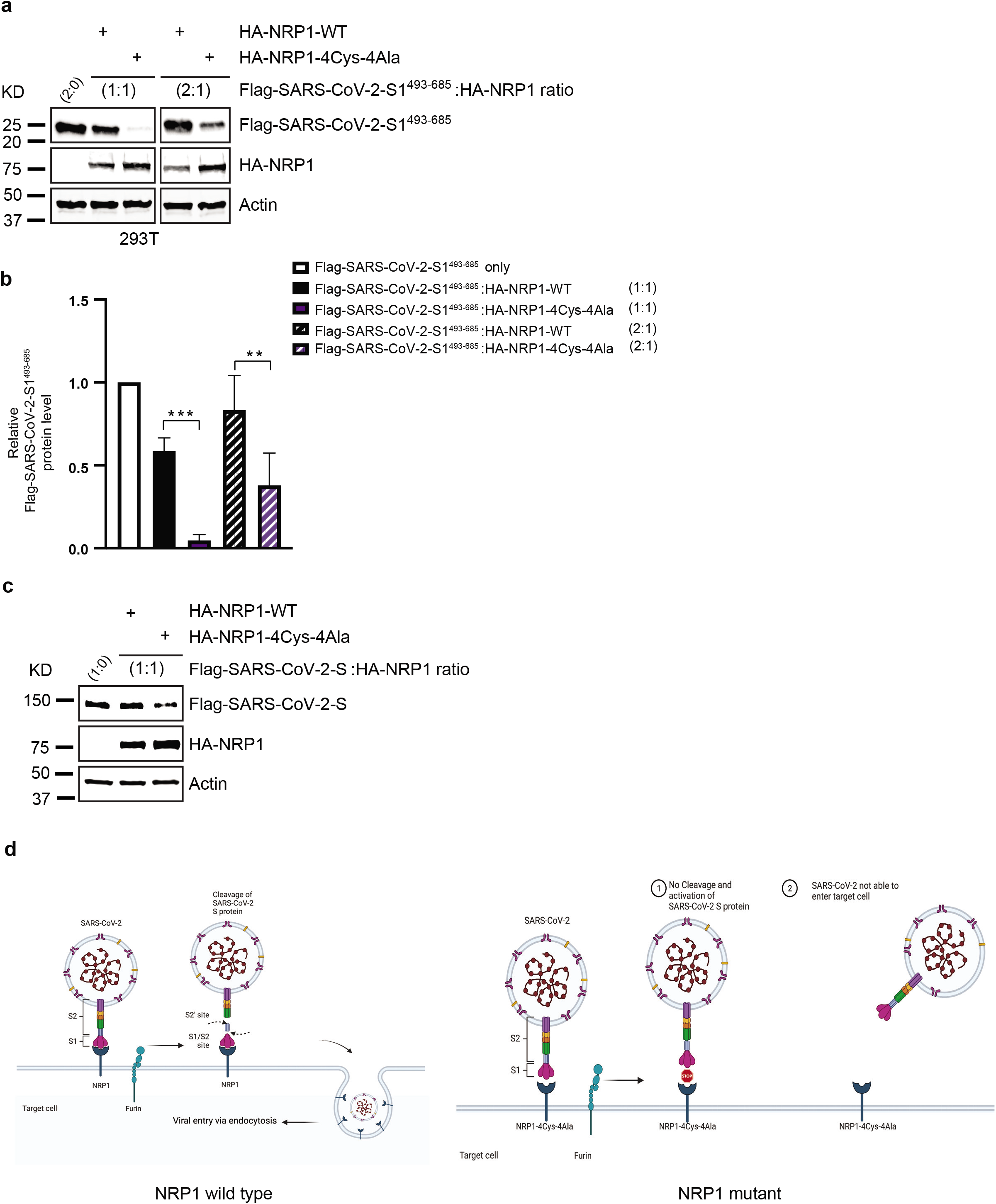
Reduced expression of SARS-CoV-2 spike protein is related to NRP1 functional mutation. (a) 293T cells were transfected with the indicated HA-NRP1 (wild type and 4Cys-4Ala mutant) and Flag-SARS-CoV-2-S1^493-685^ constructs at different ratio. Cells were lysed 30 hr post transfection, and the protein expression was analyzed. (b) Quantification of the band intensities (*n* = 3). Flag-SARS-CoV-2-S1^493-685^ band intensities were normalized to the actin band and further normalized to Flag-SARS-CoV-2-S1^493-685^ level in presence of NRP1-WT for each experimental set up. Mean and SD are indicated. Data were analyzed by one-way ANOVA: *p<0.05, **p < 0.005, *** p < 0.0005. (c) 293T cells were transfected with the indicated HA-NRP1 (wild type and 4Cys-4Ala mutant) and Flag-SARS-CoV-2-S constructs at 1:1 ratio. Cells were lysed 30 hr post transfection, and the protein expression was analyzed. (d) A proposed model showing how novel cysteines in NRP1 extracellular domains determine the fate of SARS-CoV-2 virus entry. (left panel) Intact novel cysteines in NRP1 determine its functional capability by providing the catalytic core to SARS-CoV-2 spike protein leading towards the internalization of virus in the host cells. (right panel) Alteration to those cysteine residues significantly reduce the binding between viral spike protein and NRP1 and blocks the virus entry into the host cells. Figure created with BioRender.com.

An important strategy to fight against viral infections is to identify endogenous host factors used by viral machinery for fostering their infection and replication. One way to address this question is to assess transcriptional responses of cell in response to infection. To that end, we characterized transcriptional responses in cells co-transfected either with HA-NRP1-WT and Flag-SARS-CoV-2-S1^493-685^ or HA-NRP1-4Cys-4Ala and Flag-SARS-CoV-2-S1^493-685^. Total RNA was extracted from HEK293T cells 30 hrs post-transfection. Transcriptional profiling captured significantly altered expression for several viral-infection marker genes (Fig. S3a). Specifically, we observed reduced expression for multiple ribosomal proteins in cells transfected with HA-NRP1-4Cys-4Ala. A majority of these are known to play critical role in the life cycle of viruses by promoting viral infection(*25*). For instance, it has been reported that RPL3 mutation leads to rapid loss of the M1 virus(*26*). We also observed reduced expression of RPL13 in cells transfected with mutated HA-NRP1-4Cys-4Ala. RPL13 overexpression promotes the induction and activation of promoters for the nuclear factor-κB (NF-κB) and interferon-β (IFN-β) genes(*27*). The NF-κB pathway is already considered a potential therapeutic target to treat COVID-19 disease(*28*). Ribosomal proteins RPLP1 and RPLP2 (RPLP1/2) are among the most crucial putative host factors required for the progression of dengue viruses (DENV), yellow fever virus (YFV), and Zika virus (ZIKV) infection(*29*). Therefore, reduced expression of these might be important in diminishing SARS-CoV-2 infection. RPS9(*30*), and RPLP1/2 are also required for viral protein synthesis(*29*). Human immunodeficiency virus type 1 (HIV-1) Tat, an essential regulator of viral transcription, inhibits cell proliferation through its interaction with RPS3, which also exhibited significantly reduced expression in our experiment(*31*). Interestingly, TGFB1-Induced Anti-Apoptotic Factor 1 (TIAF1) had reduced expression in cells transfected with HA-NRP1-4Cys-4Ala and has been previously shown to be involved in HPV16 infection(*32*). Another notable gene with reduced expression was DNAJB1/HSP40 cellular heat shock protein 40 (Hsp40/DnaJB1) which facilitates nuclear import of incoming influenza A virus (IAV) viral ribonucleoproteins (vRNPs) and is vital for efficient IAV replication(*33*). Human cytomegalovirus and host Transforming Acidic Coiled-Coil Containing Protein (TACC3) controls the spreading of infection in host cells(*34*). TACC3 had reduced expression in this study as well. Cumulatively, the reduced expression of genes known to be involved in viral infection and replication is consistent with results presented above. This supports our conclusion that these cysteine residues are essential for SARS-CoV-2 infection.

Additionally, we evaluated expression patterns of genes that participate in cancer pathways in our transcriptome analysis since cancer patients have significantly increased risk from COVID-19 infection(*35*). Overexpression of NRP1 has been shown to increase tumor growth, invasiveness and is associated with poor prognosis in tumors from the gastrointestinal (GI) tract, prostate, lung, ovary, as well as gliomas, osteosarcomas and melanomas(*36–39*). The use of NRP1 antagonists are well known in cancer therapy(*40*). Besides finding decreased expression of genes involved in viral replication, we found considerably low expression of some cancer markers in cells transfected with HA-NRP1-4Cys-4Ala compared to the cells transfected with HA-NRP1-WT in the presence of SARS-CoV-2-S1^493-685^ (Fig. S3a). These included kinesin family member 20A (KIF20A), SRY-box 2 (SOX2) and proteasome subunit beta type proteins (PSMB 3/4/5/6/7). KIF20A, which was initially discovered as a key protein in mitosis, was later found to be responsible for poor prognosis in cancers when it is upregulated(*11*). SOX2 is a universal cancer stem cell marker, accountable for the maintenance of cancer stem cells and tissue homeostasis(*41*), while PSMB3, 4, and 5 are cancer markers for glioma(*42*). Growth of lung cancer has been reported to be inhibited by targeting PSMB6(*43*). PSMB7 is a prognostic biomarker for breast cancer(*44*).

On the other hand, transcriptome profiling revealed upregulated expression of several genes implicated in cellular pathways like ubiquitination, degradation and suppression of inflammation. We observed significantly higher expression of both Degradation in ER protein 3 (DERL3) and Homocysteine inducible ER protein with ubiquitin like domain 1 (HERPUD1), both of which are known to promote degradation of misfolded proteins in the endoplasmic reticulum (ER)(*45, 46*). Interestingly, DERL3 and has been identified as a potential candidate gene that might be regulated via SARS-CoV-2 induced DNA methylation changes COVID-19 infection(*47*). HERPUD1 which usually activates innate immune response, is an ER stress marker and was found to be upregulated in L cells infected with mouse hepatitis virus (MHV) or SARS-CoV(*48*). To our surprise, we saw higher endoplasmic reticulum oxidoreductase 1 beta (ERO1B) expression in cells transfected with HA-NRP1-4Cys-4Ala. ERO1B re-oxidizes PDIA4 leading to reactive oxygen species production(*49*). Inflammatory responses define the outcome of viral infection and hyperinflammation in COVID-19 contributes to disease severity(*50*). One of the important factors that play a central role in the initiation and resolution of this response is the NF-κB/REL family of transcription factors. NF-κB subunit c-Rel acts as a transcriptional repressor of inflammatory genes(*51*). Our results indicated higher expression for both c-REL and NF-κB1 with HA-NRP1-4Cys-4Ala in the presence of SARS-CoV-S1^493-685^.

Finally, figure. S3b shows overlap of genes found to be differentially expressed when cells were transfected with NRP1-WT vs. NRP1-4Cys-4Ala without the spike protein as well as when cells were transfected with NRP1-WT plus SARS-CoV-2-S1^493-685^ vs. NRP1-4Cys-4Ala plus SARS-CoV-2-S1^493-685^. Interestingly, the inclusion of SARS-CoV-2-S1^493-685^ dramatically increased differential expression of unique genes to 10 times higher between the two groups (Fig. S3b). Collectively, these data potentially revealed previously unknown downstream targets involved in NRP1 mediation of COVID-19 infections.

## Discussion

In this study, we report the discovery of novel cysteine residues residing in the extracellular domain of NRP1 which, when mutated, interfere with its stability as well as its ability to bind with the SARS-CoV-2 S protein. We demonstrated up to 4 times reduction in overall spike protein levels even in the presence of an intact endogenous ACE2 receptor following mutation of these residues. There was no significant association between the NRP1-4Cys-4Ala and SARS-CoV-2 variants, suggesting that these novel cysteines are critical determinants for NRP1 activity. Moreover, NRP1 has been connected to the occurrence of various cancers. Interaction between NRP1 and its ligand, VEGF_165_ known to cause pathological angiogenesis(*52*). Considered to be a therapeutic target against cancer, NRP1 signaling could be modulated by targeting these novel cysteines since our results showed cysteine-mutated NRP1 was unable to interact with its co-receptor Plexin-A1. In conclusion, we provide a model where mutation of these cysteine residues in NRP1, especially the first three located in two CUB and b1 domains, are sufficient to reduce SARS-CoV-2 spike protein entry into cells (Fig.4d).

## Materials and Methods

### PAN domain sequence alignment

Proteins with PAN domains were identified from Uniprot and were selected from 2 model organisms(*53*). The PAN domain coordinates from Uniprot were used to extract the PAN domain sequences from the full-length proteins which were then aligned with MAFFT linsi(*54*). The alignment was visualized with Geneious.

### Mammalian cell culture, transfection, and drug treatment

HeLa and HEK 293T cells were obtained from ATCC and maintained in a humidified atmosphere at 5% CO2 in Dulbecco’s Modified Eagle’s (DMEM) complete medium (Corning) supplemented with 10% fetal bovine serum (FBS; Seradigm) in 37°C. Plasmid transfections were done with TransIT-LT1 (Mirus Bio) per the manufacturer’s instructions.

### Plasmids and recombinant proteins

HA-NRP1 and different mutants of HA-NRP1, Flag-Plexin-A1, Flag-SARS-CoV-2-S-original (NC_045512.2), Flag-SARS-CoV-2-B.1.1.7-S, Flag-SARS-CoV-B.1.351-S were all obtained from GenScript (Cloned into pcDNA 3.1 vector). Flag-SARS-CoV-2-S1^493-685^ was made in GenScript.

### Immunofluorescence and confocal microscopy

HeLa cells were seeded on coverslips in 24 well plates. Where indicated, cells were transfected with HA-NRP1 WT and different PAN domain mutants together with different SARS-CoV-2 spike protein constructs as indicated for 30 hours, followed by fixation in 4% paraformaldehyde. Next, the cells were permeabilized with 0.5% Triton X-100 in PBS, washed and then blocked for 30 minutes at room temperature with 5% BSA in PBS. Cells were incubated with primary antibodies in 5% BSA in 1X-PBS with 0.5% Triton X-100 for 1 hour at room temperature. After washing the cells were incubated with appropriate secondary antibodies in 5% BSA in PBST for 30 minutes at room temperature. DNA was counterstained with 1 μg/mL Hoechst 33342 and mounted with Fluorimount G (Southern Biotech). Cells were imaged using a Zeiss LSM 710 confocal microscope.

### Antibodies

The following commercial antibodies, and the indicated concentrations, were used in this study. HA antibody (HA.C5 #18181; 1:1000) and LAMP1 antibody (#24170) were purchased from Abcam. Flag (#2368S; 1:1000). M2 anti Flag Mouse antibody (#SLBT7654; 1:5000) and Actin (#087M4850; 1:10,000) were purchased from Sigma. HA (#902302; 1:1000) antibody was purchased from Biolegend. Chemiluminescence detection was performed according to the manufacturer’s instructions (Amersham ECL Western Blotting Detection Reagent kit) followed by exposure using Chemidoc gel-documentation system (BioRad). For imaging using Li-Cor, secondary antibodies for western blotting were purchased from LI-COR Biosciences.

### Western Blotting and immunoprecipitation

For immunoprecipitation, either HA-tagged NRP1 (or mutants) and Flag-tagged SARS-CoV-2-S (or S1^493-685^) mutant or HA-tagged NRP1 (or mutants) and Flag-Plexin-A1 were expressed where indicated in 293T cells for 30 hours. Cell extracts were generated in EBC buffer, 50mM Tris (pH 8.0), 120mM NaCl, 0.5% NP40, 1mM DTT, and protease and phosphatase inhibitors tablets (Thermo Fisher Scientific). Equal amounts of cell lysates were incubated with the indicated antibodies conjugated to protein G beads (Invitrogen) respectively from 4h to overnight at 4° C. The beads were then washed with EBC buffer including inhibitors. Immunoprecipitation samples or equal amount of whole cell lysates were resolved by SDS-PAGE, transferred to PVDF membranes (Milipore) probed with the indicated antibodies, and visualized with either LiCor Odyssey infra-red imaging system or Chemiluminescence detection. For cycloheximide stability assay, 293T cells were transfected with the constructs encoding HA-NRP1 or HA-NRP1 mutants. 24 hr post transfection, cells were splitted and replated. 20 hr post splitting, cells were treated with 100 ug cycloheximide. Cells washed with PBS and lysed. Briefly, Cell extracts were generated on ice in EBC buffer, 50mM Tris (pH 8.0), 120mM NaCl, 0.5% NP40, 1mM DTT, and protease and phosphatase inhibitors tablets (Thermo Fisher Scientific). Extracted proteins were quantified using the PierceTM BCA Protein assay kit (Thermo Fisher). Proteins were separated by SDS acrylamide gel electrophoresis and transferred to IMMOBILON-FL 26 PVDF membrane (Milipore) probed with the indicated antibodies and visualized either by chemiluminescence (according to the manufacturer’s instruction) or using LiCor Odyssey infra-red imaging system.

### RNA-seq analysis

For RNA-seq analysis, Total RNA was extracted from HEK293T cell line 30 hours post transfection (HA-NRP1 and SARS-CoV-2-S1^493-685^ and or HA-NRP1-4Cys-4Ala and SARS-CoV-2-S1^493-685^) using TRIzol reagent (Invitrogen) according to the manufacturer’s instructions. Quantification and quality control of isolated RNA was performed by measuring absorbance at 260 nm and 280 nm on a NANODROP ONEC spectrophotometer (Thermo Scientific, USA). The RNA-seq run was performed with four biological replicates. Library prep and sequencing was performed by BGI using the DNBSEQ-G400 platform which generated 100bp paired-end reads. The raw RNA-seq reads have been deposited at NCBI under BioProject ID PRJNA767890. Clean reads were aligned to the human reference genome GRCh38. Reads were mapped with bowtie2 v2.2.5. (*55*). Differential expression analysis was performed with DESeq2, genes with an adjusted p-value less than 0.05 were considered differentially expressed (*56*).

### Statistical analysis

Statistical analyses were performed on individual experiments, as indicated, with GraphPad Prism 8 Software using an unpaired t-Test, equal variance for comparison between two groups and one-way ANOVA for comparisons between more than two groups. A P value of *P<0.05 was considered as statistically significant.

## Supporting information

Figure legends for fig S1-S3

## Acknowledgements

Bioinformatics analyses of PAN domain distribution and functional inference was supported by the United States Department of Energy’s Office of Science Early Career Research Program under the Biological and Environmental Research office. Biochemical, immunofluorescence and transcriptome analyses in human cell lines was supported by the Oak Ridge National Laboratory Lab Directed Research Development program. Part of this research used resources at the Oak Ridge Leadership Computing Facility (OLCF) and the Compute and Data Environment for Science (CADES) at the Oak Ridge National Laboratory. Oak Ridge National Laboratory is managed by UT-Battelle, LLC for the U.S. Department of Energy under Contract Number DE-AC05-00OR22725.

## Author contributions

Conceptualization: DP, KD, WM

Methodology: DP and KD did all biochemical and immunofluorescence studies. KD and DP did the RNA-seq data analysis. TBY did the alignment and RNA-seq data submission to NCBI.

Funding acquisition: WM

Writing – original draft: DP, KD, WM

Writing – review & editing: DP, KD, TBY, WM

## Data and materials availability

All data are available in the main text or the supplementary materials.

