## Supplementary figures and images for "Mutating novel interaction sites in NRP1 reduces SARS-CoV-2 spike protein internalization"

### Supplemental fig 1

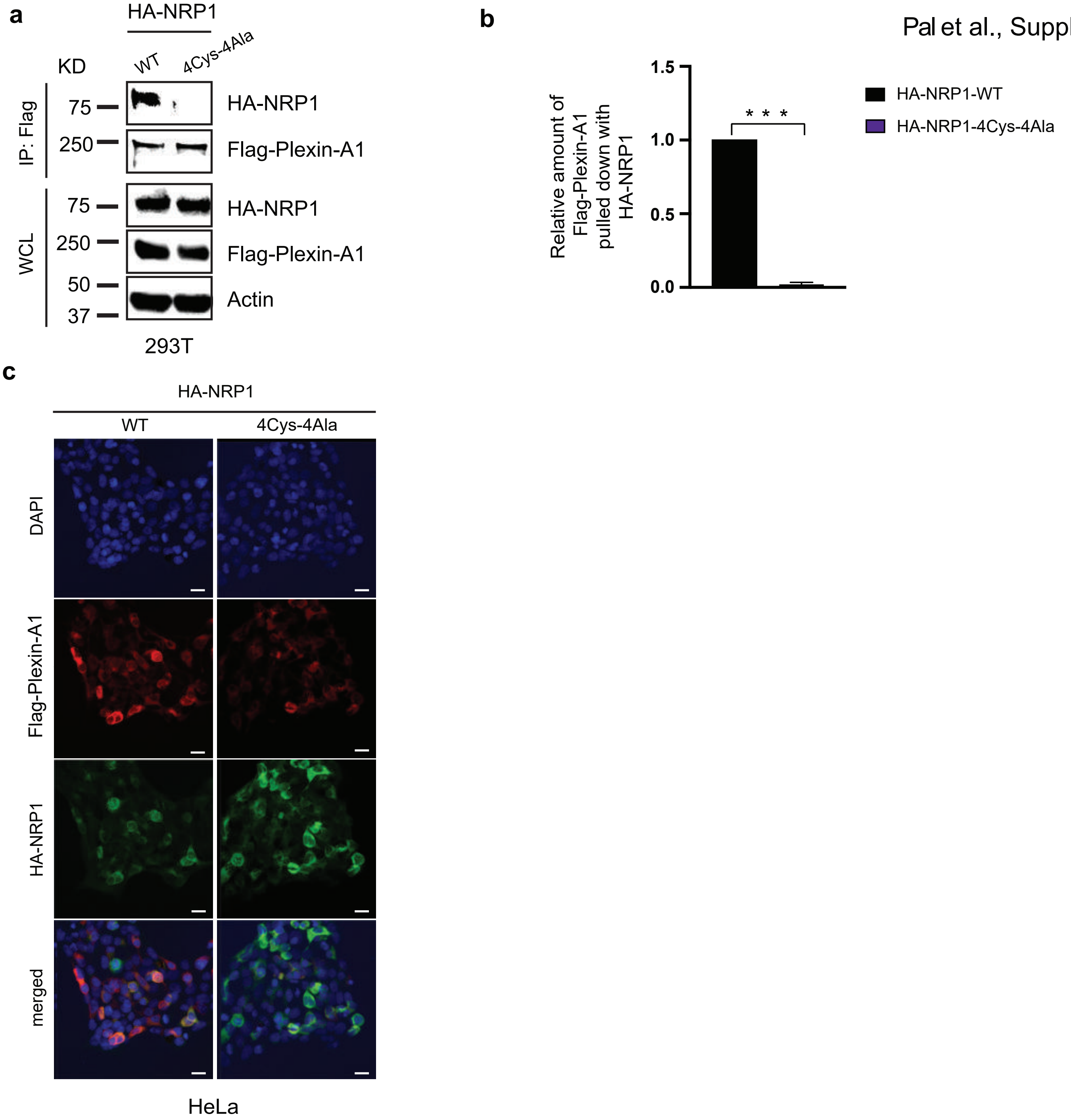

### Supplemental fig 2

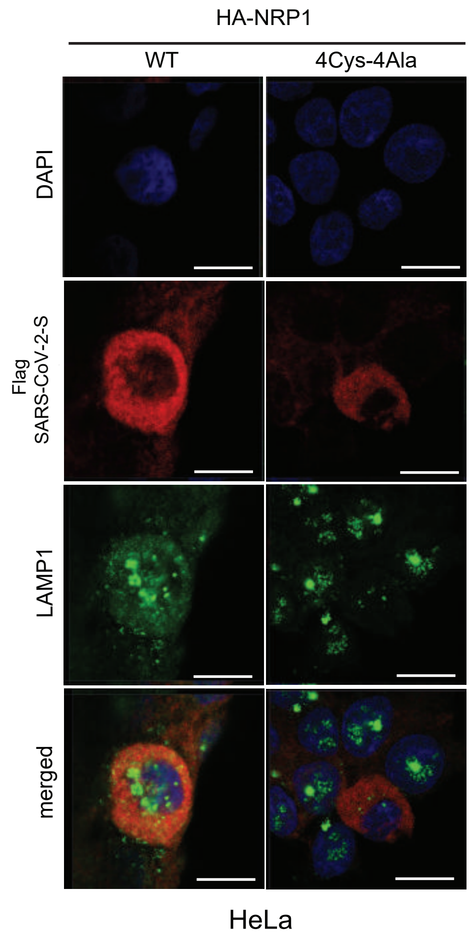

### Supplemental fig 3

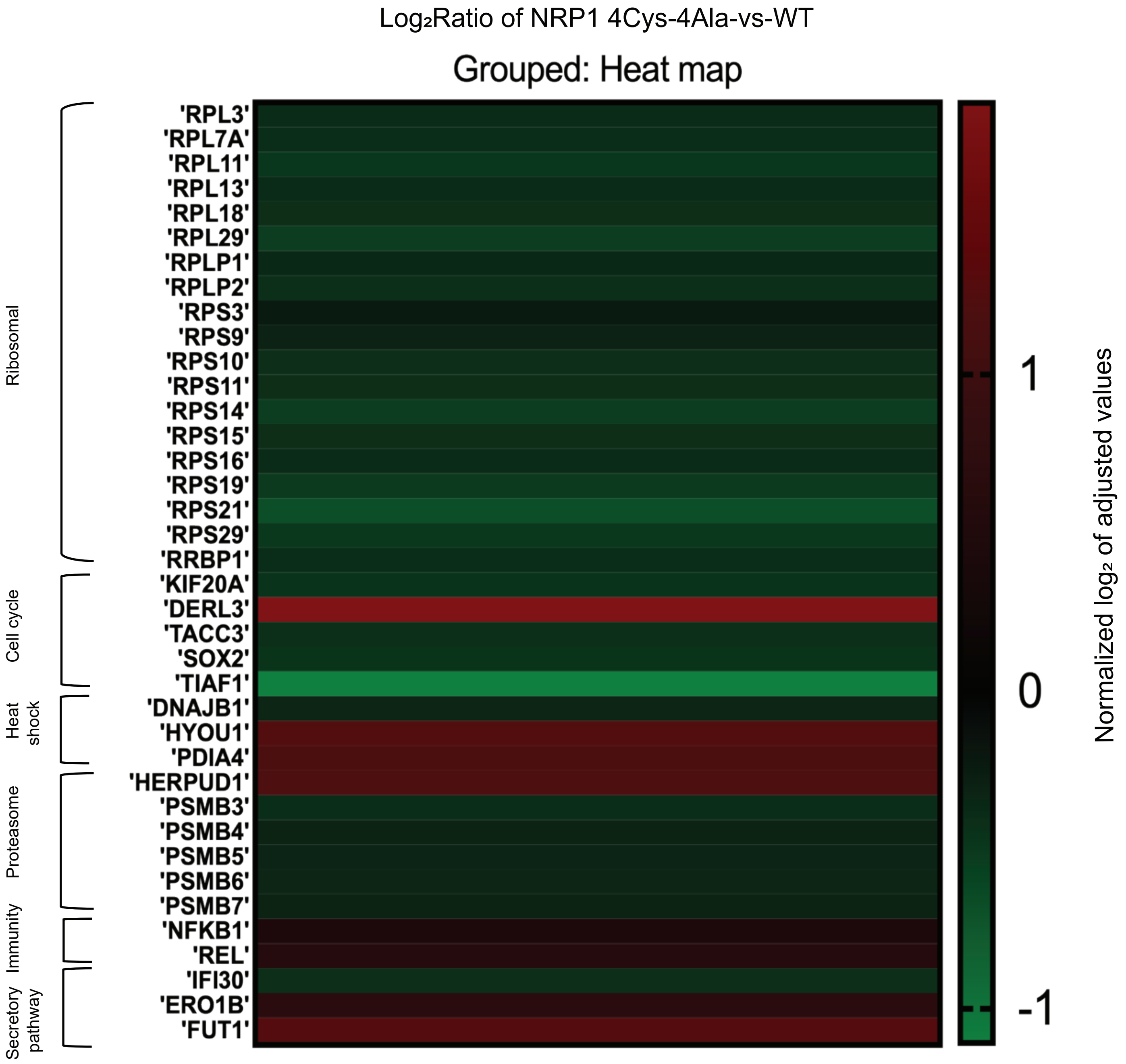

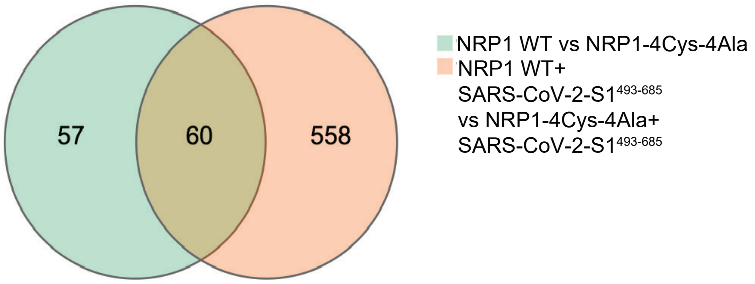
