## Supplementary material for "Mutating novel interaction sites in NRP1 reduces SARS-CoV-2 spike protein internalization": Figure legends for fig S1-S3

**This PDF file includes:**

Figs. S1 to S3

Legends for Figs.S1 to S3

**Fig. S1: NRP1 interaction with co-receptor Plexin-A1 is cysteine dependent.** (a) 293T cells were transfected with HA-NRP1-WT or HA-NRP1-4Cys-4Ala together with Flag-Plexin-A1. Cell lysates were then immunoprecipitated with anti-Flag antibody and blotted with anti-HA or anti-Flag antibody. (b) Quantification of the band intensities ($n = 2$). Immunoprecipitated HA-NRP1 band intensities were normalized to the respective Flag-Plexin-A1 IP bands and then further normalized to HA-NRP1-WT control. Data are represented as mean ± SD, and *** $p < 0.0005$ (Student's t test). (c) Representative images of colocalization studies between Flag-tagged Plexin-A1 protein and indicated HA-NRP1 constructs by confocal immunofluorescence microscopy in HeLa cells. The cells were transiently transfected with Flag-Plexin-A1 and different constructs of HA-NRP1 as indicated. 30 hours post-transfection cells were fixed, mounted and protein expression patterns were visualized using a Zeiss LSM 710 confocal microscope outfitted with a 63x objective. Scale bars represent 20 μm. The images shown are representative from three independent biological experiments (average 100 cells were observed per experimental condition per replicate).

**Fig. S2: Altered association of SARS-CoV-2 spike protein with lysosomal marker LAMP1 in the presence of mutant NRP1.** Representative image of HeLa cells transfected together with Flag-SARS-CoV-2 spike protein and different HA-NRP1 constructs (both WT and 4Cys-4Ala) and then coimmunostained with DAPI (blue), antibody specific to spike protein (Flag, red) and anti-LAMP1 (green). Cells were visualized using a Zeiss LSM 710 confocal microscope outfitted with a 63x objective. Scale bars represent 5 μm.

**Fig. S3: Transcriptome analysis in 293T cells:** (a) Heatmap of selected genes. Heatmap displaying pattern of expression for the candidates showed an altered expression pattern following transfection with different NRP1 constructs (WT and 4Cys-4Ala) and Flag-SARS-CoV-2-S1$^{493-685}$. Genes are depicted based on their expression ratios across three RNA seq comparison. Colors range from bright red (upregulation; $log_2$ ratio over control) to bright green (downregulation; $log_2$ ratio over control). (b) Venn diagram is to visualize the overlap in the genes found to be differentially expressed in the NRP1-WT vs NRP1- 4Cys-4Ala mutant and NRP1-WT+ SARS-CoV-2-S1$^{493-685}$ vs NRP1- 4Cys-4Ala+ SARS-CoV-2-S1$^{493-685}$. Each circle represents a group of gene sets, and the areas superimposed by different circles represent the intersection of these gene sets. The non-overlapping part indicate the uniquely expressed genes, and the numbers on the figure indicate the number of genes in the corresponding area.
